## Supplementary material for "Preparatory attentional templates in prefrontal and sensory cortex encode target-associated information": https://osf.io/xw8hm/

**Supplementary Materials**

| Table S1. Whole-brain searchlight results of brain regions showing significant decoding during the ***search*** ***cue*** period. | | | | |
| --- | --- | --- | --- | --- |
| Brain region | L/R | Cluster size (voxel number) | Peak MNI coordinate (x,y,z) | Z-score |
| ***Decoding of face information*** |  |  |  |  |
| Fusiform gyrus | R | 616 | 42 -48 -8 | 4.74 |
| Middle frontal gyrus | L | 2039 | -22 32 30 | 4.21 |
| Middle frontal gyrus | R | 1170 | 28 30 34 | 3.86 |
| Inferior parietal sulcus | R | 690 | 38 -48 52 | 3.82 |
| Superior parietal lobule | R |  | 30 -50 54 | 3.72 |
| Middle temporal gyrus | R | 220 | 42 -46 18 | 3.26 |
| Middle occipital gyrus | L | 182 | -14 -96 -6 | 3.67 |
| Calcarine | L |  | -20 -94 2 | 3.31 |
| Calcarine | R | 182 | 20 -92 2 | 3.39 |
| Inferior occipital gyrus | R |  | 32 -78 -8 | 3.38 |
| Precuneus | L | 424 | -10 -52 38 | 3.58 |
| ***Decoding of scene information*** |  |  |  |  |
| Insula | R | 385 | 38 12 0 | 3.91 |
| Inferior frontal gyrus | R |  | 36 24 14 | 3.66 |
| Inferior frontal gyrus | L | 268 | -38 30 28 | 3.50 |
| Middle frontal gyrus | L |  | -34 34 16 | 3.44 |
| Peak voxel coordinate is defined in MNI152 standard space. Voxel size: 2.0 2.0 2.0 mm mm mm; L, left; R, right; MNI, Montreal Neurological Institute. | | | | |

| Table S2. Whole-brain searchlight results of brain regions showing significant decoding during the ***search*** ***delay*** period. | | | | |
| --- | --- | --- | --- | --- |
| Brain region | L/R | Cluster size (voxel number) | Peak MNI coordinate (x,y,z) | Z-score |
| ***Decoding of face information*** |  |  |  |  |
| None |  |  |  |  |
| ***Decoding of scene information*** |  |  |  |  |
| Retrosplenial cortex | L | 3065 | -14 -54 12 | 4.73 |
| Retrosplenial cortex | R | 914 | 14 -48 12 | 3.98 |
| Precentral | R | 491 | 50 2 42 | 3.75 |
| Inferior frontal gyrus | R |  | 48 6 32 | 3.54 |
| Precentral | L | 192 | -42 -2 32 | 2.83 |
| Precentral | L | 54 | -36 -22 54 | 2.20 |
| Superior frontal gyrus | R | 605 | 18 36 38 | 3.08 |
| Precuneus | R | 205 | 10 -60 46 | 2.45 |
| Parahippocampal | L | 83 | -32 -36 -16 | 2.34 |
| Parahippocampal | R | 165 | 28 -58 -12 | 2.17 |
| Peak voxel coordinate is defined in MNI152 standard space. Voxel size: 2.0 2.0 2.0 mm mm mm; L, left; R, right; MNI, Montreal Neurological Institute. | | | | |

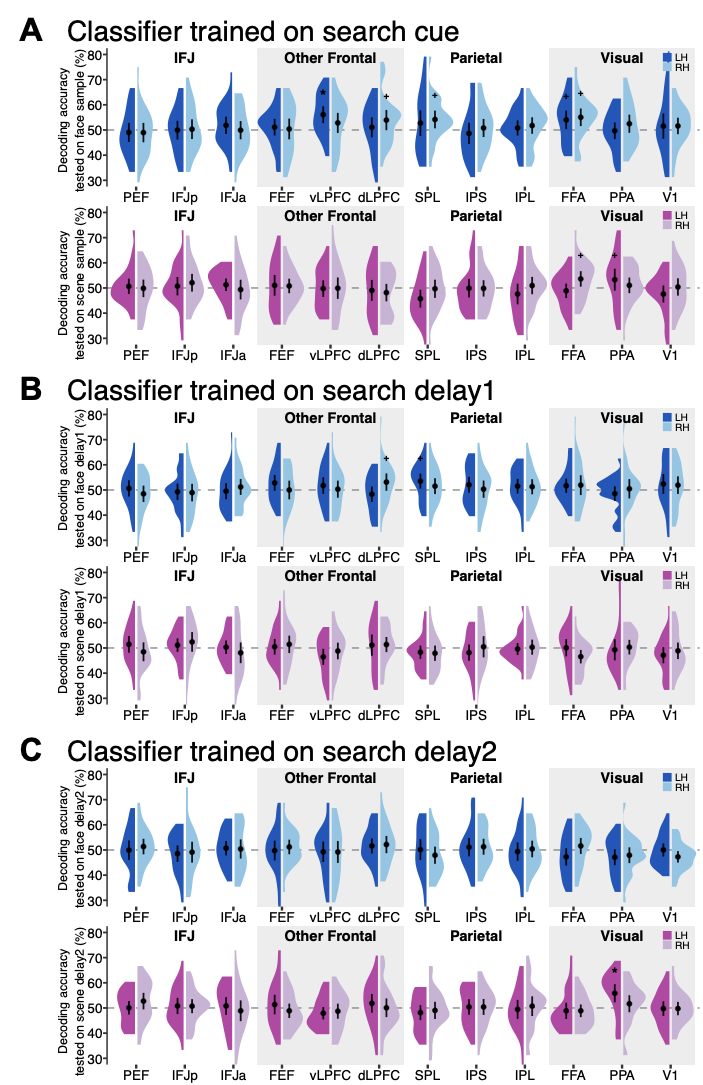

**Figure S1.** Decoding of face and scene information during the *search cue* (A), *search delay1* (B)*, and search delay2* (C) periods. An exploratory analysis was conducted to examine possible differences in decoding of target-associated information in the earlier versus later portions of the delay period. New GLMs were run. They were identical to the main analysis, except that the delay period was split into two 4-second regressors (i.e., the 1st and 2nd half of the 8 s delay period) in both the *face search task* and the *face and scene 1-back task*. The decoding schemes were identical to the main analysis but were now conducted separately for the *delay1* and *delay2* time periods. The ROI decoding results were similar for target faces during the *search cue* period using this new GLM compared to the main analysis results – this was expected since there were no differences in the model for the *search cue* period. However, decoding of scenes and faces during the *delay1* and *delay2* periods were not reliably found based on the new GLMs. The only significant effect was in LH PPA during *delay2*. The null results could be due to insufficient power when the data are divided, individual differences in when preparatory activation is the strongest, or truly no difference in activation over the delay period. Other methods with higher temporal resolution may be better suited to answer the question of exactly when preparatory activation for target-associated information is initiated and how long it lasts.

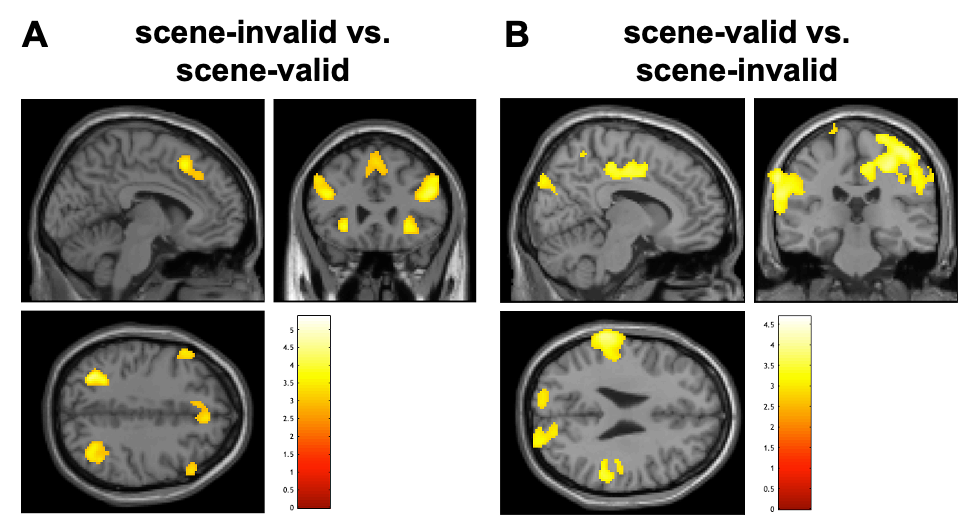

**Figure S2.** Univariate contrast results from the search period shown in a volumetric MNI standard brain. (A) Scene-invalid minus scene-valid trials. (B) Scene-valid minus scene-invalid trials. Both activation maps are shown with correction at the cluster level, *p*TFCE < .005.

| Table S3. Univariate contrasts related to scene-validity during the search period. | | | | |
| --- | --- | --- | --- | --- |
| Brain region | L/R | Cluster size (voxel number) | Peak MNI coordinate (x,y,z) | Z-score |
| ***Contrast: scene-invalid vs. scene-valid*** | | | | |
| Inferior frontal gyrus | L | 832 | -40 8 28 | 4.34 |
| Inferior frontal gyrus | R | 1075 | 42 8 28 | 3.70 |
| Inferior parietal sulcus | L | 318 | -30 -64 40 | 3.68 |
| Inferior parietal sulcus | R | 561 | 36 -62 34 | 3.27 |
| Insula | L | 88 | -28 26 -4 | 3.36 |
| Insula | R | 296 | 30 32 -8 | 3.53 |
| Anterior cingulate cortex (ACC) | R | 582 | 6 40 40 | 3.04 |
| ***Contrast: scene-valid vs. scene-invalid*** | | | | |
| Superior parietal lobule | L | 1512 | -24 -44 74 | 3.93 |
| Supramarginal | L | 2980 | -62 -34 32 | 3.61 |
| Superior parietal lobule | R | 6044 | 26 -40 54 | 3.69 |
| Supramarginal | R |  | 64 -36 42 | 3.47 |
| Middle occipital gyrus | L | 1465 | -38 -86 6 | 3.19 |
| Inferior occipital gyrus | R | 1517 | 42 -86 -6 | 3.10 |
| Peak voxel coordinate is defined in MNI152 standard space. Voxel size: 2.0 2.0 2.0 mm mm mm; L, left; R, right; MNI, Montreal Neurological Institute. | | | | |

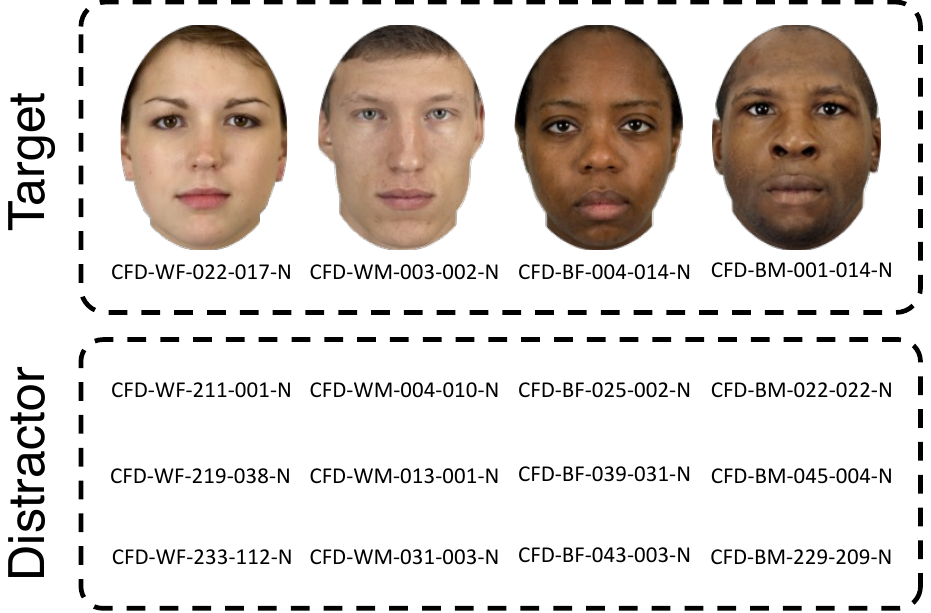

**Figure S3.** The four target face stimuli and their three distractor face counterparts. Four target faces were approved for release to the public following the Chicago Face Database (CFD; <https://www.chicagofaces.org/>) copyright rules. The image file names of the twelve distractor faces are listed for reference.

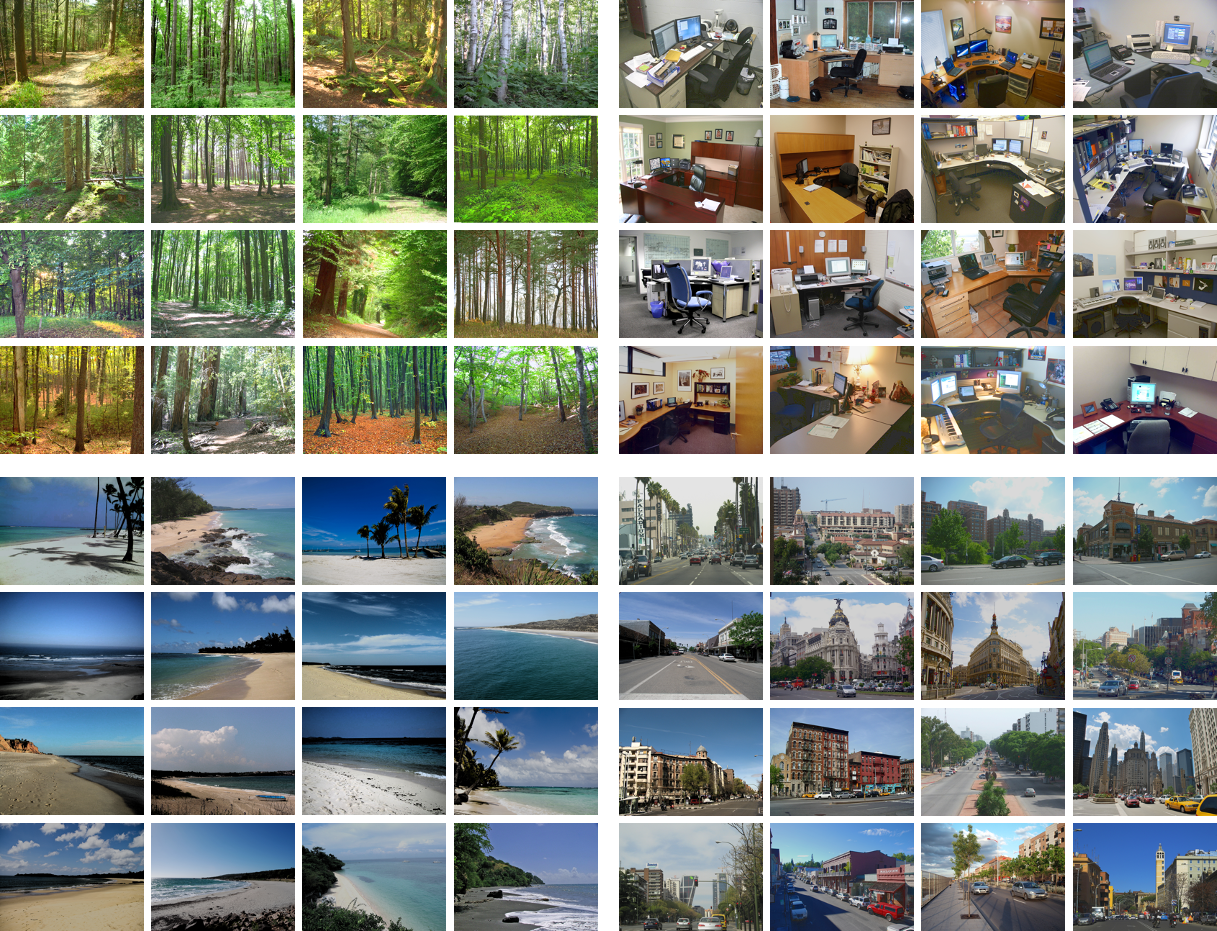

**Figure S4.** The four scene categories used in the search task. Each category consisted of 16 exemplars.

| Table S4. Mean (SE) number of voxels in each ROI | | | |
| --- | --- | --- | --- |
| ROI | | Left hemisphere | Right hemisphere |
| IFJ | PEF | 36 (5) | 47 (7) |
|  | IFJp | 29 (4) | 26 (4) |
|  | IFJa | 34 (4) | 38 (5) |
| Frontal | FEF | 83 (5) | 86 (5) |
|  | vLPFC | 269 (11) | 180 (9) |
|  | dLPFC | 44 (4) | 210 (11) |
| Parietal | SPL | 202 (7) | 205 (10) |
|  | IPS | 157 (6) | 106 (6) |
|  | IPL | 119 (5) | 155 (8) |
| Visual | FFA | 65 (6) | 113 (9) |
|  | PPA | 126 (10) | 163 (11) |
|  | V1 | 122 (7) | 175 (7) |

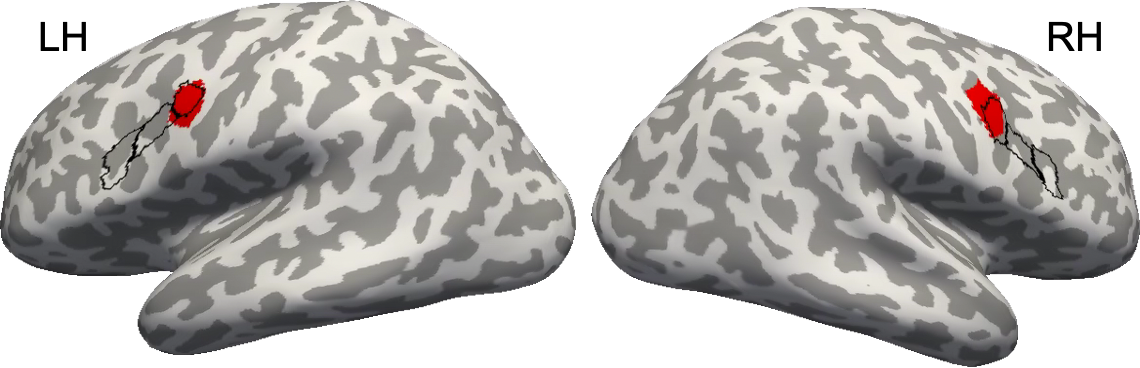

**Figure S5.** Illustration of overlap between the HCP-MMP1 atlas (Glasser et al., 2016) and the Schaefer resting state 17-network atlas (Schaefer et al., 2018) in the inferior frontal junction regions. Black outlines correspond to the PEF, IFJp, and IFJa from the HCP-MMP1 atlas. The red ROI corresponds to the 17-network atlas precentral label from the dorsal attention network in the left hemisphere and the ventral attention network in the right.
